# TRIM21–SERPINB5 aids GMPS repression to protect nasopharyngeal carcinoma cells from radiation-induced apoptosis

**DOI:** 10.1101/609743

**Authors:** Panpan Zhang, Xiaomin Li, Qiuping He, Lulu Zhang, Keqing Song, Xiaojing Yang, Qingmei He, Yaqin Wang, Xiaohong Hong, Jun Ma, Na Liu

## Abstract

Nasopharyngeal carcinoma (NPC) is the most prevalent head and neck malignancy in South China and Southeast Asia. The main NPC treatment strategy is radiotherapy. However, recurrence resulting from radioresistance is a leading clinical bottleneck. Revealing the mechanism of NPC radioresistance would help improve the therapeutic effect. Here, our work reveals that *TRIM21* (tripartite motif–containing 21) functions as an oncogene in NPC progression, and its ablation increases NPC cell radiosensitivity. Further analysis indicated that TRIM21 represses TP53 expression by mediating GMPS (guanine monophosphate synthase) ubiquitination and degradation after ionizing radiation. Mass spectrometry and co-immunoprecipitation showed that SERPINB5 (serpin family B member 5) interacts with both TRIM21 and GMPS. Epistatic analysis showed that SERPINB5 acts as an adaptor to recruit GMPS and introduce TRIM21 for ubiquitination. The in vitro and in vivo results validated the finding that SERPINB5 promotes NPC cell radioresistance. Furthermore, immunohistochemistry indicated that radioresistant patients have higher SERPINB5 expression. Overall, our data show that TRIM21–SERPINB5-mediated GMPS degradation facilitates TP53 repression, which promotes the radioresistance of NPC cells.

## Introduction

Nasopharyngeal carcinoma (NPC) is a malignant head and neck cancer with apparent regional aggregation (Jemal, Bray et al., 2011, McDermott, Dutt et al., 2001, Wei & Sham, 2005). Radiotherapy is the most effective treatment strategy against NPC. With modern intensity-modulated radiation therapy, the 5-year overall survival rate of patients with NPC is increased to nearly 80% (Wu, Liu et al., 2017). However, about 20% of patients with locoregionally advanced disease will have local or regional recurrence, and 90% of the recurrence is in the radiation field (Yi, Gao et al., 2006), owing to radioresistance of the tumor cells. Therefore, revealing the underlying mechanism governing NPC radioresistance would shed light on new clinical therapy and help improve the curative effect.

Upon DNA damage resulting from ionizing radiation or cytotoxic drugs, TP53 will activate the DNA repair system to maintain the integrity of the whole genome, while the apoptotic process will be started if the DNA damage proves irreparable. TP53-governed apoptosis is considered the main cause of ionizing radiation–induced cell death, despite the fact that some cancer cells undergo TP53-independent apoptosis (Afshar, Jelluma et al., 2006, Strasberg Rieber, Zangemeister-Wittke et al., 2001). Therefore, the radioresistant tumor cells are often accompanied by *TP53* mutation or repression, high levels of *BCL2*, or inhibition of the other apoptosis-related genes (Brown & Wouters, 1999, Leone, Humar et al., 1997, McGill & Fisher, 1997, Reed, Miyashita et al., 1996, Yaes, 1989). In NPC, it has been suggested that latent membrane protein 1 (LMP1), encoded by Epstein–Barr virus, blocks apoptosis and thereby facilitates radioresistance of the tumor cells (Lu, Ma et al., 2008). Moreover, microRNA-205 inhibits apoptosis by repressing phosphatase and tensin homolog (*PTEN*) expression in NPC (Qu, Liang et al., 2012). However, the mechanism of NPC radioresistance remains largely unknown.

*TP53* is not frequently mutated in NPC (Lo, Mok et al., 1992, Sun, Hegamyer et al., 1992). Consequently, it appears that TP53 expression and its related signaling might be suppressed in radioresistant NPC cells. The protein stability of TP53 is mainly modulated by the interplay between the ubiquitination ligase MDM2 (MDM2 proto-oncogene) and the deubiquitylating enzymes (Brooks & Gu, 2011, Frappier & Verrijzer, 2011). In normal conditions, TP53 ubiquitination and degradation sustains its low levels in the nucleus. Upon radiation or other genotoxic triggers, TP53 deubiquitylation is accelerated and the TP53 expression level increases correspondingly. Several ubiquitin-specific protease (USP) family members, including USP7 (Li, Chen et al., 2002), USP10 (Yuan, Luo et al., 2010), and USP42 (Hock, Vigneron et al., 2011), are involved in maintaining TP53 protein stability. However, how TP53 is manipulated in radioresistant NPC cells remains obscure.

Previously, our work indicated that tripartite motif–containing 21 (*TRIM21*) functions as an oncogene during NPC progression (Zhang, Li et al., 2018). Moreover, TRIM21 can repress TP53 expression by promoting guanine monophosphate synthase (GMPS) ubiquitination and degradation in genotoxic stress conditions (Reddy, van der Knaap et al., 2014). GMPS also interacts with USP7 to mediate gene transcription or H2B deubiquitylation in human cells (Frappier & Verrijzer, 2011, van der Knaap, Kozhevnikova et al., 2010, van der Knaap, Kumar et al., 2005). Whether the TRIM21–GMPS cascade is conserved in NPC and how this cascade regulates TP53 are all unclear.

Serpin family B member 5 (*SERPINB5*), also known as *MASPIN*, was first reported to function as a tumor repressor gene in breast cancer (Zou, Anisowicz et al., 1994). However, immunohistochemistry staining in a subsequent study revealed higher SERPINB5 expression levels in patients with breast cancer who had worse prognosis (Umekita, Ohi et al., 2002), complicating matters. A recent finding suggested that SERPINB5 function is determined by its cellular localization and that SERPINB5 plays a tumor suppressor role only when localized in the nucleus (Goulet, Kennette et al., 2011). In NPC, the functional mechanism of SERPINB5 is unknown.

Here, we show that TRIM21 prevented apoptosis in NPC cells after ionizing radiation by mediating GMPS ubiquitination and degradation. Mass spectrometry (MS) and co-immunoprecipitation revealed that SERPINB5 interacts with TRIM21 to facilitate GMPS repression. Moreover, SERPINB5 acts as an adaptor to recruit GMPS protein independent of TRIM21 expression, which was strengthened after radiation. NPC specimens from radioresistant patients had higher SERPINB5 expression levels than radiosensitive patients, suggesting the potential application of SERPINB5 in distinguishing radioresistant patients clinically.

## Results

First, we examined the function of TRIM21 in NPC. *TRIM21* mRNA expression levels in NPC cell lines were significantly upregulated (Fig 1A), as was that in NPC biopsy samples (Gene Series Expression [GSE]81672252, Fig 1B). Moreover, higher *TRIM21* expression levels predicted poorer overall survival in head and neck squamous cell carcinoma (HNSC) (Fig 1C). To explore the function of TRIM21 in NPC, we generated a stable *TRIM21* gain-of-function (GOF) NPC cell line, and *TRIM21* CRISPR (clustered regularly interspaced short palindromic repeats) knockout mutant (loss of function, LOF) NPC cells (Supplementary Fig 1A). TRIM21 promoted NPC cell proliferation, which was demonstrated by Cell Counting Kit-8 (CCK-8) and the colony formation assay (Fig 1D, 1E). However, there was no sign of TRIM21 involvement in NPC cell invasion (Fig 1F). Therefore, the data indicate that *TRIM21* functions as an oncogene in NPC.

**Figure 1.**
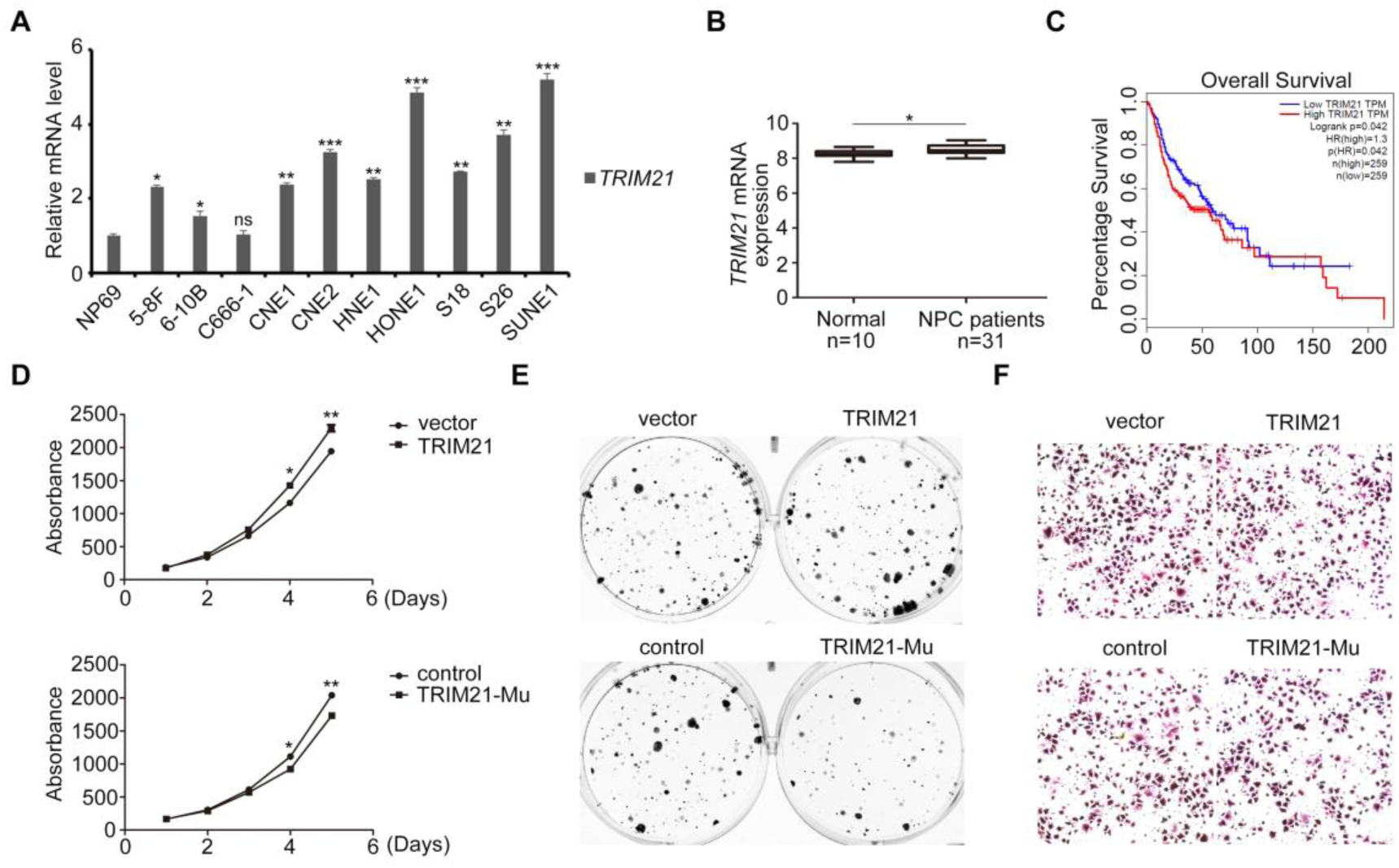
*TRIM21* serves as an oncogene in NPC. A qRT-PCR detection and comparison of *TRIM21* expression in normal NP69 NPEC line and in NPC cell lines. B *TRIM21* expression in healthy controls and patients with NPC in the Gene Expression Omnibus (GEO) dataset (GSE81672252). C Analysis of overall survival according to *TRIM21* expression level in HNSC in TCGA dataset. D CCK-8 assay of HONE1 cells with *TRIM21* GOF or LOF. E Colony formation assay of NPC cells with *TRIM21* GOF or LOF. F Transwell assay of HONE1 cells with *TRIM21* GOF or LOF. Data information: \**P* < 0.05, \*\**P* < 0.01, \*\*\**P* < 0.001. Mu, mutant; ns, not significant.

As TRIM21 protects breast cancer cells from chemotherapy-mediated apoptosis by repressing the GMPS–TP53 cascade, we wondered whether NPC cells share the same mechanism scenario after radiation. X-ray irradiation was followed by an obvious increase in TP53 expression (Fig 2A). However, this increase was reversed in NPC cells with TRIM21 ectopic expression, and vice versa (Supplementary Fig 1B). Then, we expressed FLAG-tagged GMPS in HONE1 cells, followed by anti-FLAG antibody–mediated immunoprecipitation. In the context of X-ray radiation, GMPS bound both USP7 and TP53 (Fig 2B), and TP53 protein levels were elevated upon GMPS overexpression (Supplementary Fig 1C), suggesting that GMPS promotes TP53 protein stability. In addition, TRIM21 downregulated GMPS, especially under the condition of radiation (Fig 2C).

**Figure 2.**
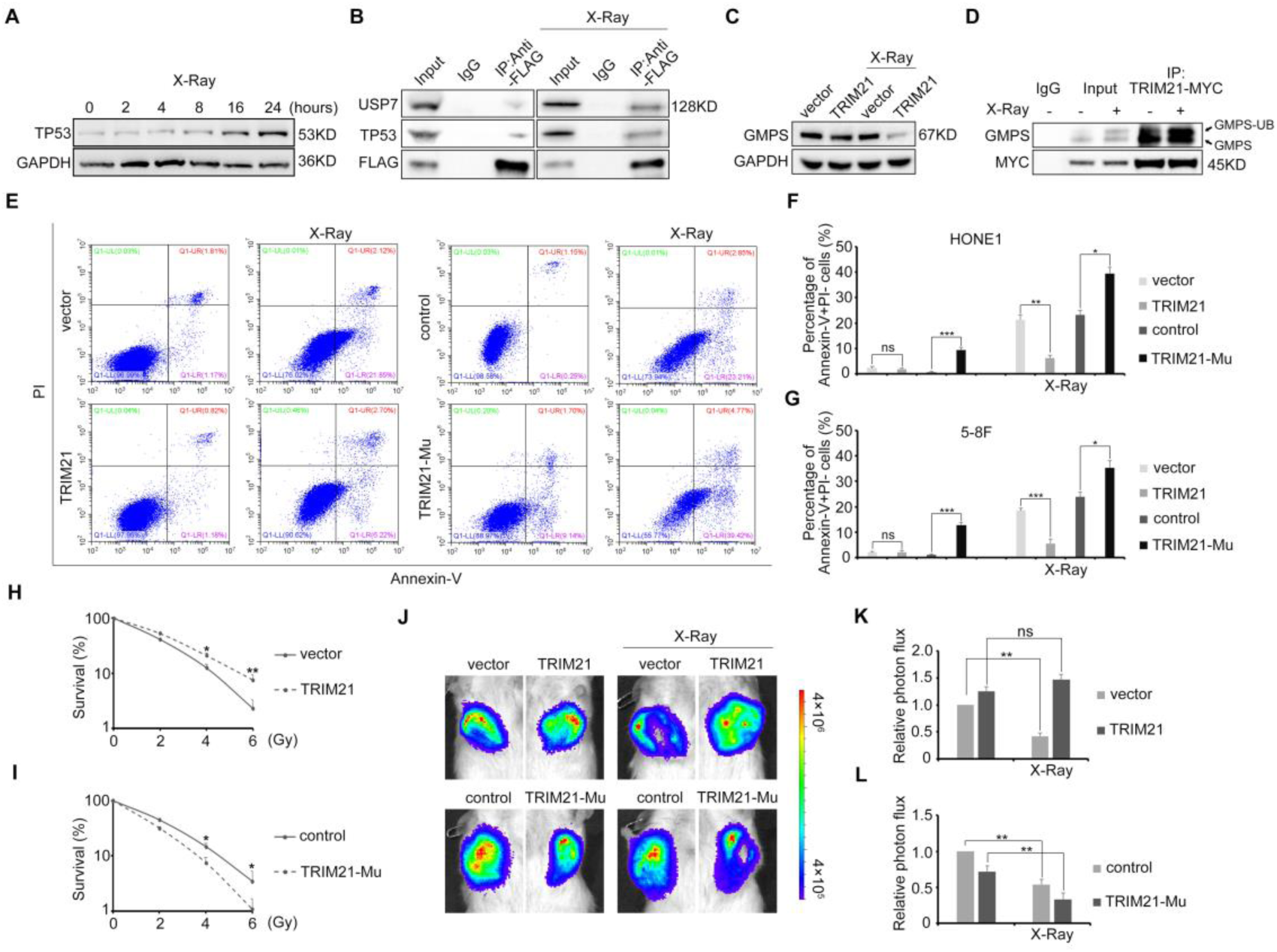
TRIM21 protects NPC cells from radiation-induced apoptosis by manipulating the GMPS–TP53 cascade. A Western blot detection of TP53 expression in HONE1 cells after radiation. B Co-immunoprecipitation following western blot detection of USP7 and GMPS in GMPS–FLAG-overexpressing NPC cells with or without X-ray radiation. C GMPS expression in TRIM21-overexpressing NPC cells with or without X-ray radiation. D Co-immunoprecipitation following western blot detection of GMPS in TRIM21–MYC-overexpressing NPC cells with or without X-ray radiation. E Flow cytometry analysis of annexin V and PI staining in HONE1 cells with *TRIM21* GOF or LOF after X-ray radiation. F, G Quantification of the apoptotic HONE1 (F) and 5-8F (G) cells. H, I Clonogenic survival assay of HONE1 cells with *TRIM21* GOF (H) or LOF (I). J Absorbance intensity of *TRIM21* GOF and LOF tumor cells and their counterpart controls in mice. The tumors were evaluated 2 and 3 weeks, respectively, after implantation, and the mice received radiotherapy (2 Gy daily and a total of 12 Gy) after 2 weeks. K, L The absorbance intensity analysis of tumors in mice. Data information: \**P* < 0.05, \*\**P* < 0.01, \*\*\**P* < 0.001. Mu, mutant; ns, not significant; IP, immunoprecipitation.

MS and immunoprecipitation were performed using TRIM21–MYC purified cell lysate. GMPS was included in the MS analysis (Supplementary Table 1). Ubiquitinated GMPS was identified in the immunoprecipitated cell lysate with TRIM21–MYC overexpression (Fig 2D), indicating that radiation facilitated TRIM21-mediated GMPS protein ubiquitination and degradation.

Based on the above findings, we deduced that altered *TRIM21* expression disrupted NPC cell apoptosis. Therefore, HONE1 and 5-8F NPC cells with *TRIM21* GOF or LOF underwent annexin V staining and flow cytometry analysis. As expected, X-ray–irradiated *TRIM21* GOF cells had significantly attenuated early apoptosis, and vice versa (Fig 2E–G). The clonogenic survival assay showed that TRIM21 elevated the survival rate of NPC cells, while TRIM21 blockage attenuated it (Fig 2H, 2I). Moreover, TRIM21 overexpression attenuated active caspase-3 expression (Supplementary Fig 1D).

To identify whether manipulating TRIM21 expression would modify NPC cell radiosensitivity in vivo, *TRIM21* GOF or LOF HONE1 cells with luciferase activity were injected subcutaneously into immunodeficient mice. Following X-ray radiation, tumor formation was observed and evaluated. Consistent with the above results, high *TRIM21* expression levels protected the tumor cells from radiation-mediated cell death, whereas *TRIM21* knockout rendered NPC cells radiosensitive (Fig 2J–L). Therefore, our data demonstrate that TRIM21 plays an essential role in regulating NPC cell radiosensitivity.

As X-ray radiation accelerated TRIM21-mediated GMPS ubiquitination, we proposed that there are radiation-activated factors that facilitate GMPS degradation. The MS data showed that SERPINB5 was highly enriched (Supplementary Table 1). The co-immunoprecipitation showed that TRIM21 interacted with SERPINB5 in NPC cells and that radiation strengthened this interaction (Fig 3A). Moreover, immunofluorescence staining revealed that SERPINB5 mainly localized in the cytoplasm, along with the colocalized TRIM21 protein (Supplementary Fig 2A, Fig 3B).

**Figure 3.**
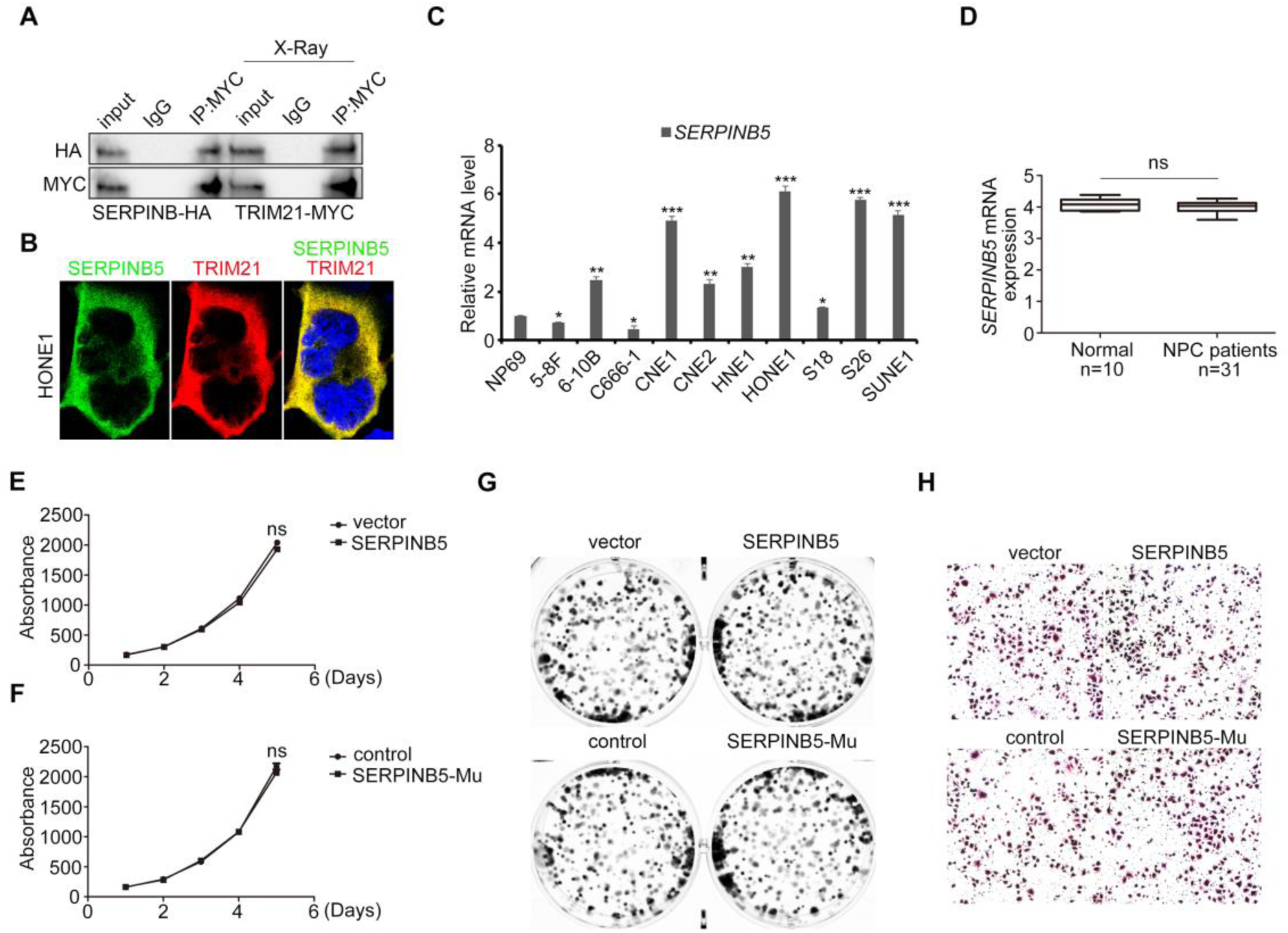
SERPINB5 does not affect NPC progression in normal conditions. A Co-immunoprecipitation following western blotting in NPC cells with SERPINB5–HA and TRIM21–MYC overexpression. B Immunofluorescence staining analysis of SERPINB5–HA and TRIM21–MYC in NPC cells. C qRT-PCR detection and comparison of *SERPINB5* expression in normal NP69 NPEC line and in NPC cell lines. D *SERPINB5* expression in healthy controls and patients with NPC in the GEO dataset (GSE81672303). E, F CCK-8 assay of NPC cells with *SERPINB5* GOF (E) or LOF (F). G Colony formation assay of NPC cells with *SERPINB5* GOF (top) or LOF (bottom). H Transwell assay of NPC cells with *SERPINB5* GOF (top) or LOF (bottom). Data information: \**P* < 0.05, \*\**P* < 0.01, \*\*\**P* < 0.001. Mu, mutant; ns, not significant; IP, immunoprecipitation.

To determine the role of SERPINB5 in NPC, we detected its mRNA level in NPC cell lines first. Surprisingly, *SERPINB5* expression was not consistent in the NPC cell lines (Fig 3C). Moreover, *SERPINB5* was not significantly different between normal and NPC biopsy samples, as well as that in HNSC (Fig 3D, Supplementary Fig 2B). To explore the function of SERPINB5 in NPC, HONE1 and 5-8F cells, which have higher and lower *SERPINB5* mRNA levels, respectively, were employed. We generated stable *SERPINB5* overexpression cells and CRISPR knockout cells (the transcription start site was mutated [Mu]) (Supplementary Fig 2C, 2D). CCK-8 and the colony formation assay revealed that SERPINB5 did not function during NPC cell proliferation (Fig 3E–G). In addition, the Transwell assay showed that SERPINB5 was not related to the cell invasive process (Fig 3H).

Then, we asked whether SERPINB5 is involved in TRIM21-mediated GMPS–TP53 repression. Western blotting indicated that TRIM21-mediated TP53 attenuation was dependent on SERPINB5 expression, even in the context of X-ray radiation (Fig 4A). Next, we examined the counteraction between SERPINB5 and GMPS. GMPS was precipitated by anti-HA (hemagglutinin) antibody, and X-ray radiation promoted the interaction (Fig 4B). Immunofluorescence indicated that ionizing radiation strengthened the GMPS and SERPINB5 colocalization (Fig 4C).

**Figure 4.**
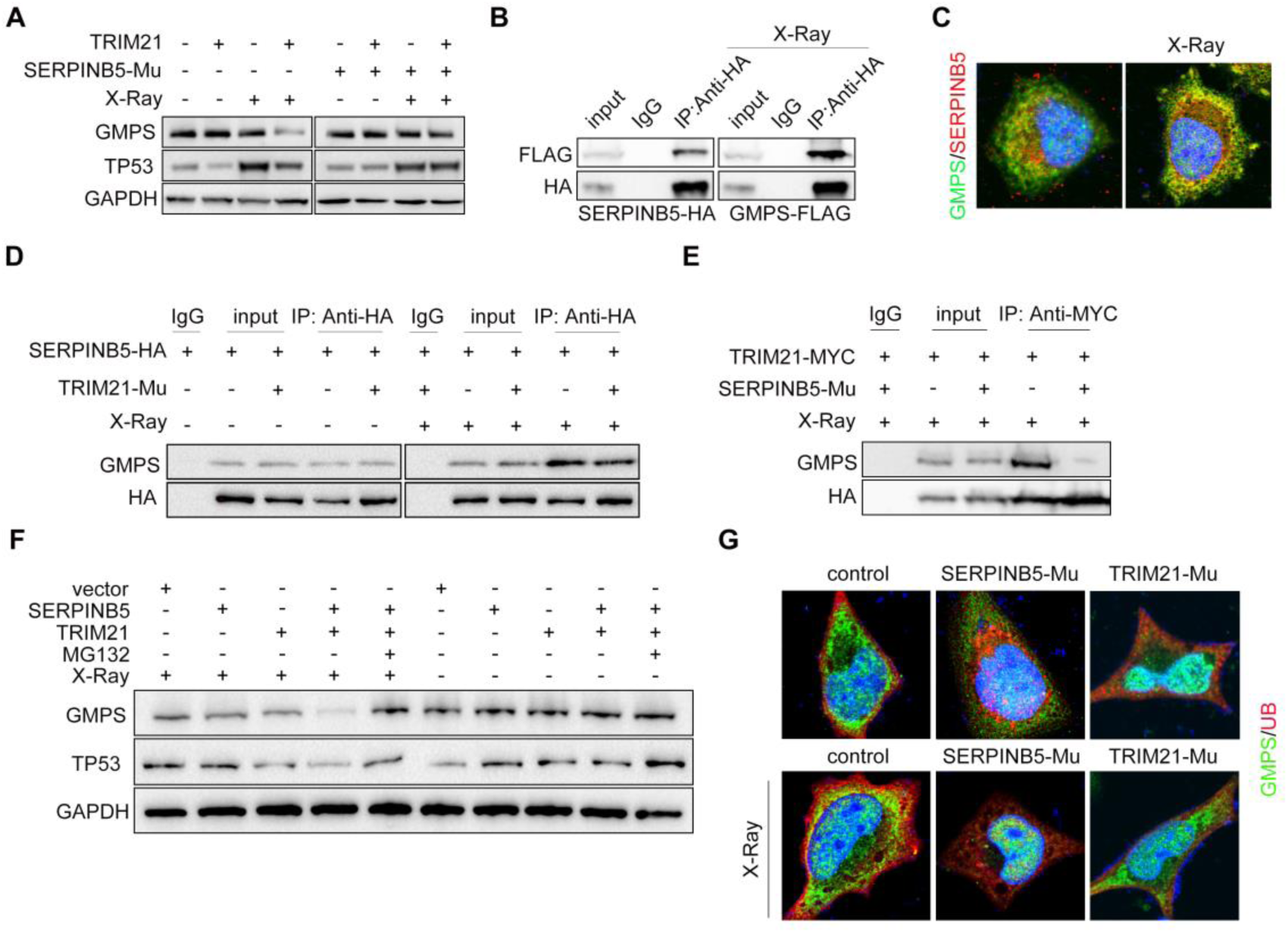
SERPINB5 is indispensable for TRIM21-mediated GMPS–TP53 repression after radiation. A Western blot detection of GMPS and TP53 in NPC cells with *TRIM21* GOF or *SERPINB5* LOF. B Co-immunoprecipitation following western blot examination of NPC cells with SERPINB5–HA and GMPS–FLAG overexpression. C Immunofluorescence staining of overexpressed GMPS and SERPINB5 in NPC cells with or without ionizing radiation. D GMPS expression in immune-purified protein from NPC cells with *SERPINB5* GOF or *TRIM21* LOF. E GMPS expression in immune-purified protein from NPC cells with *TRIM21* GOF or *SERPINB5* LOF. F GMPS and TP53 expression in NPC cells with TRIM21 or SERPINB5 overexpression. G Immunofluorescence staining of overexpressed GMPS and ubiquitin in HONE1 cells with or without ionizing radiation. Data information: Mu, mutant; IP, immunoprecipitation.

Then, we wondered whether the interaction between SERPINB5 and GMPS was dependent on TRIM21. Endogenous GMPS expression was evaluated in cell lysates with *SERPINB5* ectopic expression or *TRIM21* mutation. Our data suggested that SERPINB5 binding of GMPS was dependent on X-ray stimulation, regardless of *TRIM21* expression (Fig 4D). Correspondingly, the interaction between TRIM21 and GMPS was dependent on SERPINB5 expression, even in the condition of irradiation (Fig 4E). In addition, GMPS protein was subjected to proteasome-dependent degradation in NPC cells after the radiation, which the concomitant TRIM21 and SERPINB5 overexpression accelerated (Fig 4F).

According to previous findings, GMPS stabilizes TP53 after entering the nucleus(Reddy et al., 2014). Therefore, we examined the localization of GMPS in cells with or without ionizing radiation. GMPS localized in both the cytoplasm and the nucleus, while ionizing radiation facilitated GMPS ubiquitination in the cytoplasm, which was not observed in the *TRIM21* mutant cells (Fig 4G). Moreover, GMPS mainly localized in the nucleus in *SERPINB5* mutant cells after radiation (Fig 4G). Then, we detected GMPS expression in the cytoplasm and the nucleus. TRIM21-mediated GMPS downregulation in the cytoplasm was dependent on SERPINB5, and GMPS protein tended to localize in the nucleus without SERPINB5 (Supplementary Fig 2E). These data suggest that SERPINB5 is irreplaceable in mediating GMPS ubiquitination by TRIM21.

Next, we speculated that SERPINB5 might also play roles in governing NPC cell radiosensitivity. Annexin V staining followed by flow cytometry analysis revealed that ectopic expression of SERPINB5 protected the tumor cells from radiation-induced apoptosis, and vice versa (Fig 5A–C). The clonogenic survival assay showed that tumor cells lacking *SERPINB5* became sensitive and vulnerable to radiation (Supplementary Fig 3A, 3B). In addition, *SERPINB5* mutation completely blocked the radioresistant effect of TRIM21 (Supplementary Fig 3C, 3D), suggesting that TRIM21 acts through SERPINB5 to manipulate tumor cell radiosensitivity.

**Figure 5.**
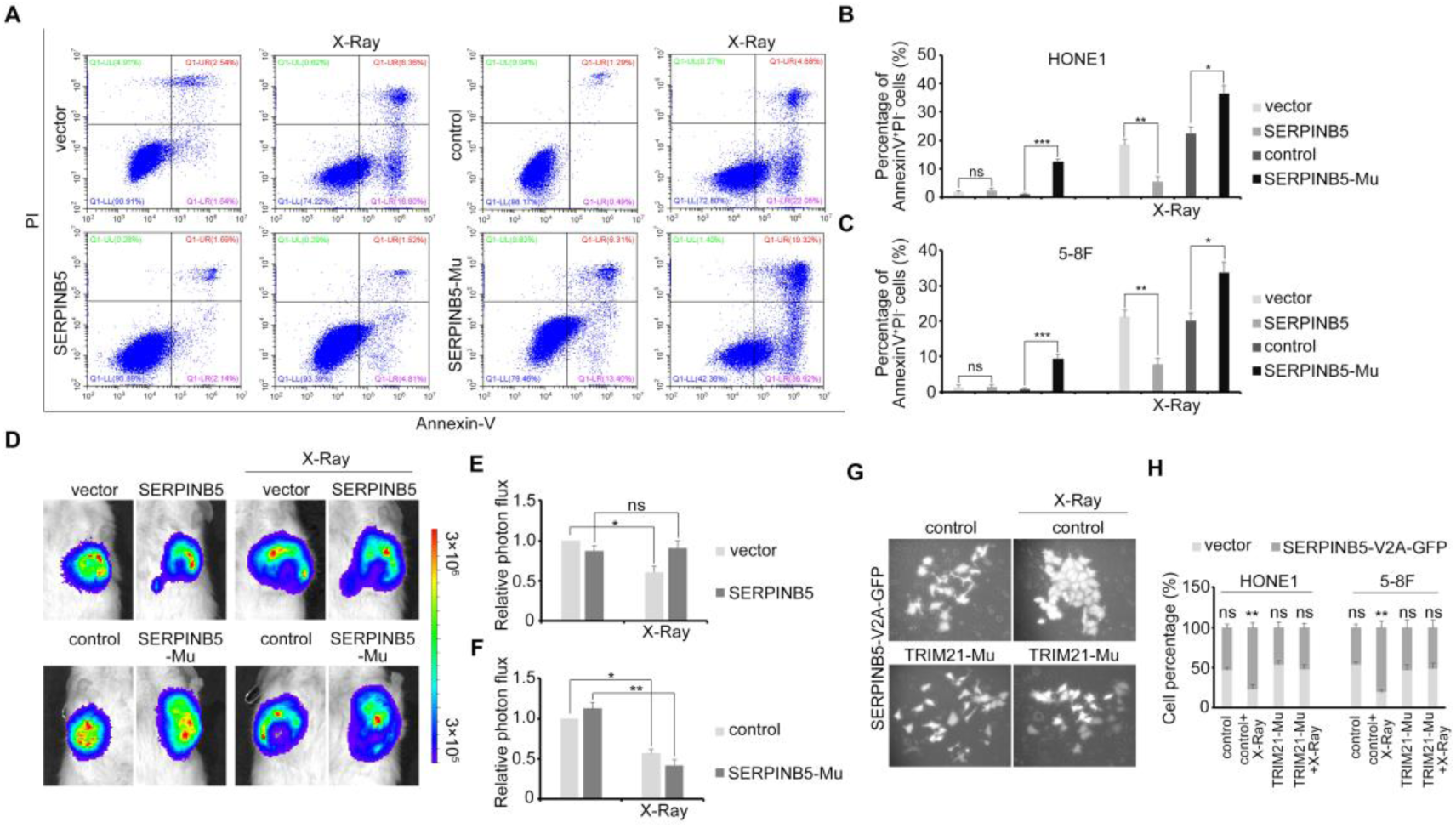
SERPINB5 prevents X-ray radiation–induced NPC cell apoptosis. A Flow cytometry analysis of annexin V and PI staining in HONE1 cells with *TRIM21* GOF or LOF after X-ray radiation. B, C Quantification of apoptotic HONE1 (B) and 5-8F (C) cells. D Absorbance intensity of *SERPINB5* GOF (top) and LOF (bottom) tumor cells and their counterpart controls in mice. The tumors were evaluated 2 and 3 weeks, respectively, after implantation, and the mice received radiotherapy (2 Gy daily and a total of 12 Gy) after 2 weeks. E, F Absorbance intensity analysis of the tumors in mice. G GFP expression of HONE1 cells with SERPINB5-V2A-GFP overexpression or TRIM21 knockout. H Flow cytometry analysis of GFP^+^ cell percentages in HONE1 and 5-8F cells. Data information: \**P* < 0.05, \*\**P* < 0.01, \*\*\**P* < 0.001. Mu, mutant; ns, not significant.

To demonstrate the effect of SERPINB5 in vivo, HONE1 cells with *SERPINB5* GOF or LOF were injected subcutaneously into immunodeficient mice, followed by regular X-ray radiation and observation. SERPINB5 rendered the tumor cells resistant and refractory to radiotherapy (Fig 5D–F). To confirm the radioresistant role of SERPINB5 in NPC, we constructed a V2A vector system with simultaneous SERPINB5 and GFP (green fluorescent protein) co-expression. The dynamic expression of GFP was evaluated after X-ray radiation in SERPINB5-V2A-GFP expression cells that had been mixed with their control cell counterparts. The percentage of GFP-positive cells increased significantly after radiation, while this increase was abrogated in the cells without *TRIM21* (Fig 5G, 5H), indicating that SERPINB5-mediated tumor cell radioresistance is dependent on TRIM21. These data demonstrate that SERPINB5 and TRIM21 function together as pivotal regulators during NPC radiotherapy.

As shown above, patients with NPC had upregulated *TRIM21* expression, while *SERPINB5* expression varied between patients. As the patients had varied outcomes after radiotherapy, we hypothesized that the SERPINB5 expression level determines the radiosensitivity of patients with NPC. To prove this, we used specimens from four radiosensitive patients and eight patients refractory to radiotherapy. Immunohistochemistry staining revealed that all radioresistant NPC samples had higher SERPINB5 expression levels (Fig 6A, 6B). Moreover, GMPS expression correlated negatively with SERPINB5 somewhat, illustrating the validity of the TRIM21–SERPINB5–GMPS signaling axis in NPC.

**Figure 6.**
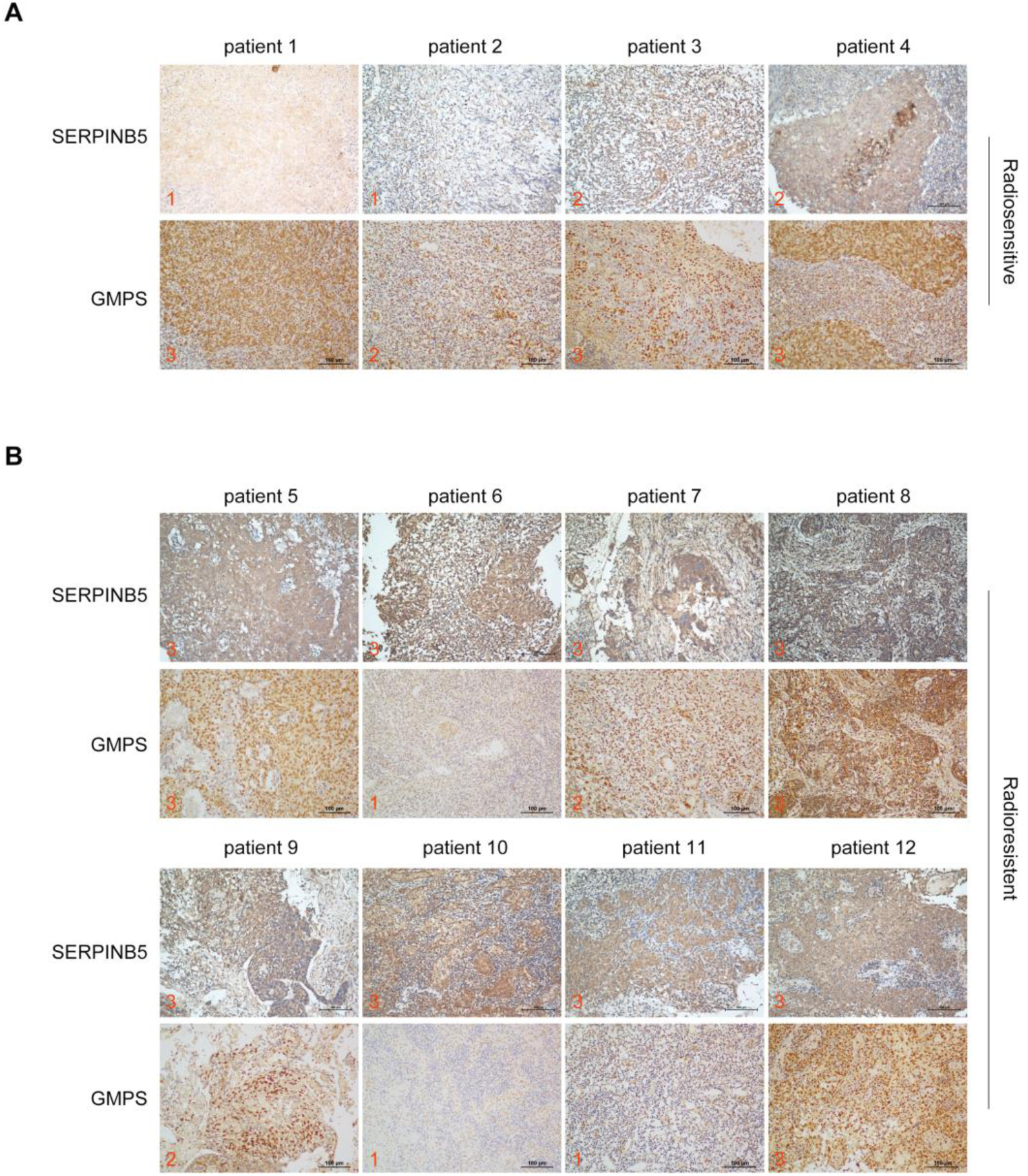
SERPINB5 expression increases in radioresistant patients with NPC. A, B Immunohistochemistry staining of SERPINB5 and GMPS in radiosensitive (A) and radioresistant (B) patients. Data information: Based on the staining intensity, the images are divided into three grades from weakest to strongest (from 1 to 3, respectively).

In general, our work reveals the regulatory mechanism of TRIM21-mediated GMPS–TP53 repression in NPC, emphasizing the critical role of SERPINB5 during this process. SERPINB5 could recruit free GMPS protein in the cytoplasm, presenting it to TRIM21 for ubiquitination and protein degradation, and X-ray radiation accelerated this process. The decreased expression of GMPS subsequently promoted TP53 degradation, which prevented radiotherapy-induced apoptosis (Supplementary Fig 4).

## Discussion

Radioresistance is one of the main obstacles in NPC clinical therapies. However, the mechanism of NPC radioresistance has remained obscure to date. In modern therapeutic strategies against NPC, all patients receive definitive radiotherapy of similar intensity (Li, Sun et al., 2012, Luo, Deng et al., 2004, Ma, Liu et al., 2007, Zhao, Han et al., 2004) despite their tumor heterogeneity. Therefore, some patients with high radiosensitivity experience adverse effects from the radiation, while some patients with low radiosensitivity face the risk of recurrence after radiation. Distinguishing the radioresistant patients is one of the main difficulties restricting improvement of the cure rate. Our findings provide a molecular marker for predicting the radiosensitivity of patients with NPC before treatment.

Our data suggest that *SERPINB5* does not influence NPC progression in normal conditions, while serving as an oncogene after radiation. We demonstrate for the first time that SERPINB5, which mainly localizes in the cytoplasm, functions as an adaptor to bind and prevent GMPS protein from entering the nucleus, and prompts GMPS ubiquitination by interacting with TRIM21. The stimulation by X-ray radiation strengthened this process in NPC. These findings stress the pivotal role of SERPINB5 in mediating GMPS–TP53 cascade repression in radioresistant NPC cells. However, how SERPINB5 detects the radiation signal remains unclear.

Our data reveal that TRIM21 promotes NPC progression in normal conditions, suggesting that it is also involved in other signaling axes in NPC. GMPS was originally found to fuel cancer progression by mediating guanine nucleotide synthesis (Su, Wiltshire et al., 2004, Weber, 1983). The Cancer Genome Atlas (TCGA) dataset showed upregulated GMPS expression in various cancers, including NPC. Considering GMPS expression was not decreased or even elevated in some of the radioresistant patients in the present study, we believe that GMPS plays multiple roles in NPC. Therefore, unlike SERPINB5, GMPS is not a suitable marker for identifying radioresistant patients with NPC.

In summary, our work establishes a novel working model related to TP53 suppression in radioresistant NPC cells, and highlights the important potential application of SERPINB5 in predicting the radiosensitivity of patients with NPC.

## Materials and Methods

### Ethical approval

This study was performed in accordance with ethical standards, according to the Declaration of Helsinki, and according to national and international guidelines. The Sun Yat-sen University Cancer Center ethics committee approved the study.

### Patients and tumor tissue samples

Tumor samples were obtained from patients with pathologically confirmed NPC (n = 12) at Sun Yat-sen University Cancer Center. Radioresistant patients were defined as those with local recurrent disease at the nasopharynx and/or neck lymph nodes at ≤12 months after completion of radiotherapy. Radiosensitive patients were defined as those without local residual lesions at >3 months and without local recurrent disease >12 months after completion of radiotherapy. All patients provided written informed consent.

### Cell lines

NP69, a human immortalized nasopharyngeal epithelial (NPEC) cell line, was cultured in keratinocyte serum-free medium (Invitrogen, Life Technologies, Grand Island, NY, USA) with bovine pituitary extract (BD Biosciences, San Jose, CA, USA). The NPC cell lines 5-8F, 6-10B, C666-1, CNE1, CNE2, HNE1, HONE1, S18, S26, and SUNE1 were cultured in RPMI 1640 medium (Invitrogen) supplemented with 5% fetal bovine serum (FBS, Gibco, Carlsbad, CA, USA). The cells were seeded in 6-well plates the day before transfection, which was performed using Lipofectamine 3000 (Invitrogen), and the cells were harvested 2 days later. For X-ray irradiation, the adhered cells received a 6-Gy dose by an X-ray irradiation apparatus (RS2000, Rad Source, Buford, GA, USA), and were harvested 24 hours later.

### RNA extraction and reverse transcription–PCR (RT-PCR)

Total RNA was extracted from the cell lines using TRIzol (Invitrogen). Complementary DNA (cDNA) was synthesized using M-MLV (Moloney murine leukemia virus) reverse transcriptase (Promega, Madison, WI, USA), and amplified using SYBR Green qRT-PCR SuperMix-UDG reagents (Invitrogen) and a CFX96 instrument (Bio-Rad, Hercules, CA, USA). The genes were amplified using the following forward and reverse primers: *GAPDH* (glyceraldehyde-3-phosphate dehydrogenase), 5′-GAAGGTGAAGGTCGGAGT-3′ and 5′-GAAGATGGTGATGGGATTTC-3′; *TRIM21*, 5′-CCCCTCTAACCCTCTGTC-3′ and 5′-GCTAAAGCTCGCTTGCTG-3′; *SERPINB5*, 5′-CATAGAGGTGCCAGGAGC-3′ and 5′-GAACAGAATTTGCCAAAGAA-3′.

### Western blot, co-immunoprecipitation, and immunofluorescence

Total protein was extracted using radioimmunoprecipitation assay lysis buffer (Beyotime, Shanghai, China). Proteins were separated by sodium dodecyl sulfate–polyacrylamide gel electrophoresis and transferred onto polyvinylidene difluoride membranes (Millipore, Billerica, MA, USA). The membranes were then incubated with primary antibodies at 4°C overnight. After incubation with species-matched secondary antibodies, immunoreactive proteins were detected using chemiluminescence in a gel imaging system (ChemiDoc MP Imaging System, Bio-Rad). The antibodies used were against the following: HA (1:2000, H6908, Sigma-Aldrich, Munich, Germany), FLAG (1:2000, F2555, Sigma-Aldrich), MYC (1:2000, 60003-2-Ig, Proteintech, Chicago, IL, USA), TRIM21 (1:1000, 12108-1-AP, Proteintech), SERPINB5 (1:1000, ab182785, Abcam, Cambridge, MA, USA), GMPS (1:1000, 16376-1-AP, Proteintech), GAPDH (1:2000, ab8245, Abcam), TP53 (1:1000, ab26, Abcam), caspase-3 (1:2000, ab32351, Abcam), lamin B1 (1:1000, ab16048, Abcam), and USP7 (1:5000, 66514-1-Ig, Proteintech).

For co-immunoprecipitation, cells with ectopic expression of SERPINB5, TRIM21, or GMPS were cultured with MG132 (10 μM, S2619, Selleck Chemicals, Houston, TX, USA) to inhibit proteasome-mediated protein degradation. After 24 hours, the cells were harvested for protein purification. The protein was incubated with the corresponding tag antibodies at 4°C overnight, followed by 3–4-hour incubation at 4°C with protein A/G agarose (20421, Invitrogen). The beads were then collected for western blot detection. The antibodies used in the co-immunoprecipitation were against the following: HA (1:100, H6908, Sigma-Aldrich), MYC (1:100, 60003-2-Ig, Proteintech), FLAG (1:100, F2555, Sigma-Aldrich), and immunoglobin G (IgG, 1:100, sc-398703, Santa Cruz, Dallas, TX, USA).

For immunofluorescence, 1×10 ^5^ cells overexpressing SERPINB5, GMPS, or TRIM21 were seeded and cultured on cover glass, and fixed with methanol after 24 hours. The cells were then incubated with tag antibodies at 4°C overnight, followed by reaction with the corresponding secondary fluorescent antibody (1:500, A-21206, A-21203, Invitrogen) and Hoechst staining (1:5000, H3570, Invitrogen). Images were captured using confocal microscopy (Olympus, Tokyo, Japan).

### Mass spectrometry

HONE1 cells with TRIM21 ectopic expression were harvested for immunoprecipitation. Then, the purified protein underwent MS analysis by Huijun Biotechnology (Guangzhou, China). The enriched protein was obtained by comparing with the IgG group.

### Flow cytometry

For apoptosis analysis, 1 × 10^5^ cells were seeded in 6-well plates. Before the cell density reached about 90%, the cells were collected and stained with annexin V and propidium iodide (PI, C1062, Beyotime), and analyzed by flow cytometry (CytoFLEX 1, Beckman Coulter, Brea, CA, USA). For GFP percentage analysis, 5 × 10^4^ cells with SERPINB5-V2A-GFP overexpression were mixed with 5 × 10^4^ vector control cells and seeded in 6-well plates. Before the cell density reached about 70%, the cells were treated with or without X-ray irradiation, and were harvested 48 hours later for GFP analysis.

### Stable cell line establishment and CRISPR gene knockout

The *TRIM21, SERPINB5*, and *GMPS* coding sequences were cloned separately into pSin-EF2-puro vector. Stable overexpression cell lines were obtained by puromycin screening and confirmed by western blotting. For CRISPR-mediated gene knockout, the genomic RNAs (gRNAs) were searched (https://zlab.bio/guide-design-resources) and cloned into lentiCRISPRv2 vector. The constructs were transfected into NPC cells, followed by puromycin screening. The surviving cells were confirmed by western blotting. For the single-clone surviving cells, the gDNA was extracted for mutation site identification. The gRNA sequences are as follows. *TRIM21*, 5′-AGCACGCTTGACAATGATGT-3′, *SERPINB5*, 5′-AGCCGAATTTGCTAGTTGCA-3′, and the *SERPINB5* forward and reverse verification primer sequences were 5′-ACTGGGCTCCCGACAATG-3′ and 5′-GCAGGCTGAGGCACAACA-3′, respectively.

### Cell proliferation, colony formation, and cell invasion assays

CCK-8 was used to detect cell proliferative ability. Cells (1 × 10^3^) were seeded in 96-well plates, incubated for 0–4 days, and stained using CCK-8 (Dojindo, Tokyo, Japan). The absorbance was determined at 450 nm using a spectrophotometer.

For the colony formation assay, about 300 cells were seeded in 6-well plates. After 7–10-day culture, the cells were fixed in methanol and stained with crystal violet.

For the cell invasion assay, 3 × 10^3^ cells were seeded in 24-well Transwell chambers (Corning, NY, USA). The medium was supplemented with 10% FBS and placed in the bottom chambers. After 14–18-hour culture, the chambers were collected and the cells on lower surface of the chambers were fixed in methanol and stained with crystal violet for observation.

### Clonogenic survival assay

The clonogenic survival assay was performed as previously reported (Munshi, Hobbs et al., 2005). HONE1 or 5-8F cells were harvested after receiving X-ray radiation, and were re-seeded in 6-well plates and incubated for 12–14 days. Then, the cell colonies were stained with crystal violet and counted. The survival rate of each group was calculated according to the corresponding plating efficiency.

### Animal experiments

B-NDG mice (non-obese diabetes, severe combined immunodeficiency with double knockout of the interleukin-2 receptor gamma chain and protein kinase DNA-activated catalytic genes: NOD-*Prkdc*^*scid*^ *IL2rg*^*tm1*^/Bcgen) were purchased from Biocytogen Jiangsu Co., Ltd. (Jiangsu, China). The cell groups were all transfected with CMV-luciferase plasmid, and about 1 × 10^6^ cells were injected subcutaneously into the dorsal or ventral flank. The mice were monitored after 7–10 days. Luciferin was diluted to 15 mg/ml using phosphate-buffered saline, and 100 μl of the solution was injected intraperitoneally into each mouse. After 5 minutes, the mice were anesthetized and observed using an animal imaging system (IVIS Lumina LT, PerkinElmer, Waltham, MA, USA). All animal research was performed in accordance with the detailed rules approved by the Sun Yat-sen University Cancer Center Animal Care and Use Ethics Committee; all efforts were made to minimize animal suffering.

### Immunohistochemistry

Paraffin-embedded patient samples were sectioned and mounted on slides. The slides were incubated at 4°C overnight with antibody against SERPINB5 (1:200, ab182785, Abcam) or GMPS (1:100, 16376-1-AP, Proteintech). Then, the sections were incubated with biotinylated secondary antibody bound to a horseradish peroxidase complex. The antibody was visualized by adding 3,3-diaminobenzidine, and the sections were counterstained with hematoxylin.

### Statistical analysis

Statistical analyses were performed using SPSS 17.0 (SPSS Inc., Chicago, IL, USA). All in vitro data shown are representative of at least three independent experiments, and values are reported as the mean ± SD. Differences between two groups were analyzed using the two-tailed unpaired Student’s *t*-test; *P* < 0.05 was considered significant.

## Acknowledgments

This work was supported by grants from the National Natural Science Foundation of China (81802920), Natural Science Foundation of Guangdong Province (2017A030310227, 2018B030306045, 2017A030312003), Guangdong Special Support Program (2017TQ04R754), and The Health & Medical Collaborative Innovation Project of Guangzhou City, China (201803040003). The funders had no role in the study design, data collection, analysis, decision to publish, or the preparation of the manuscript.

## Authors’ contributions

PP.Z., XM.L, and QP.H. carried out all experiments, prepared the figures, and drafted the manuscript. LL.Z., KQ.S., XJ.Y., QM.H., YQ.W., and XH.H. participated in the data analysis and interpretation of results. PP.Z., J.M., and N.L. conceived the study and participated in the data analysis. All authors read and approved the manuscript.

---

**Supplementary Figure 1.**
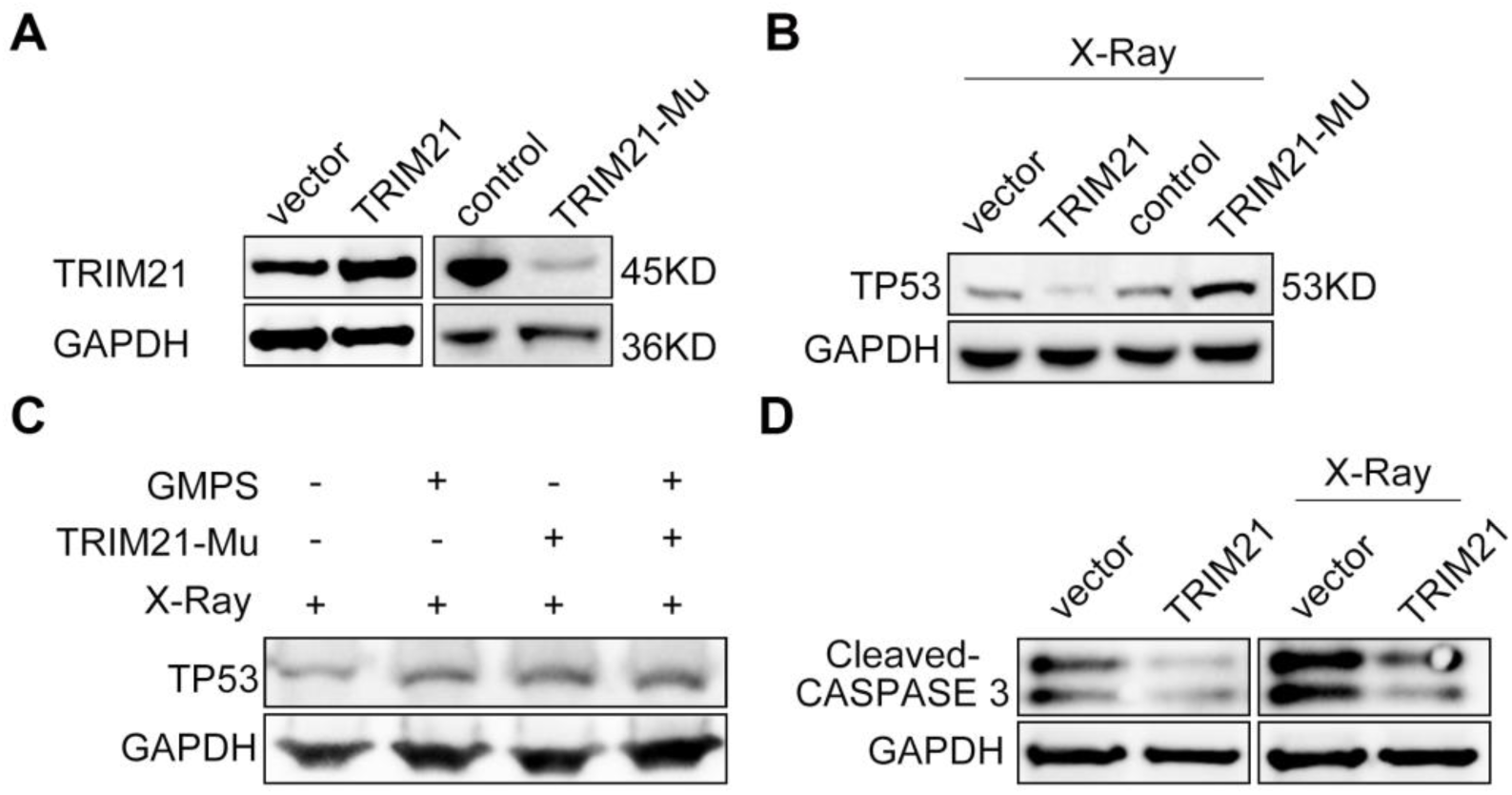
The TRIM21–GMPS signaling cascade regulated TP53. A Western blot detection of TRIM21 expression in *TRIM21* knockout HONE1 cells. B TP53 expression in NPC cells with *TRIM21* GOF or LOF after X-ray radiation. C TP53 expression in NPC cells with *GMPS* GOF or *TRIM21* LOF after X-ray radiation. D Cleaved caspase-3 expression in HONE1 cells with *TRIM21* overexpression. Data information: Mu, mutant.

**Supplementary Figure 2.**
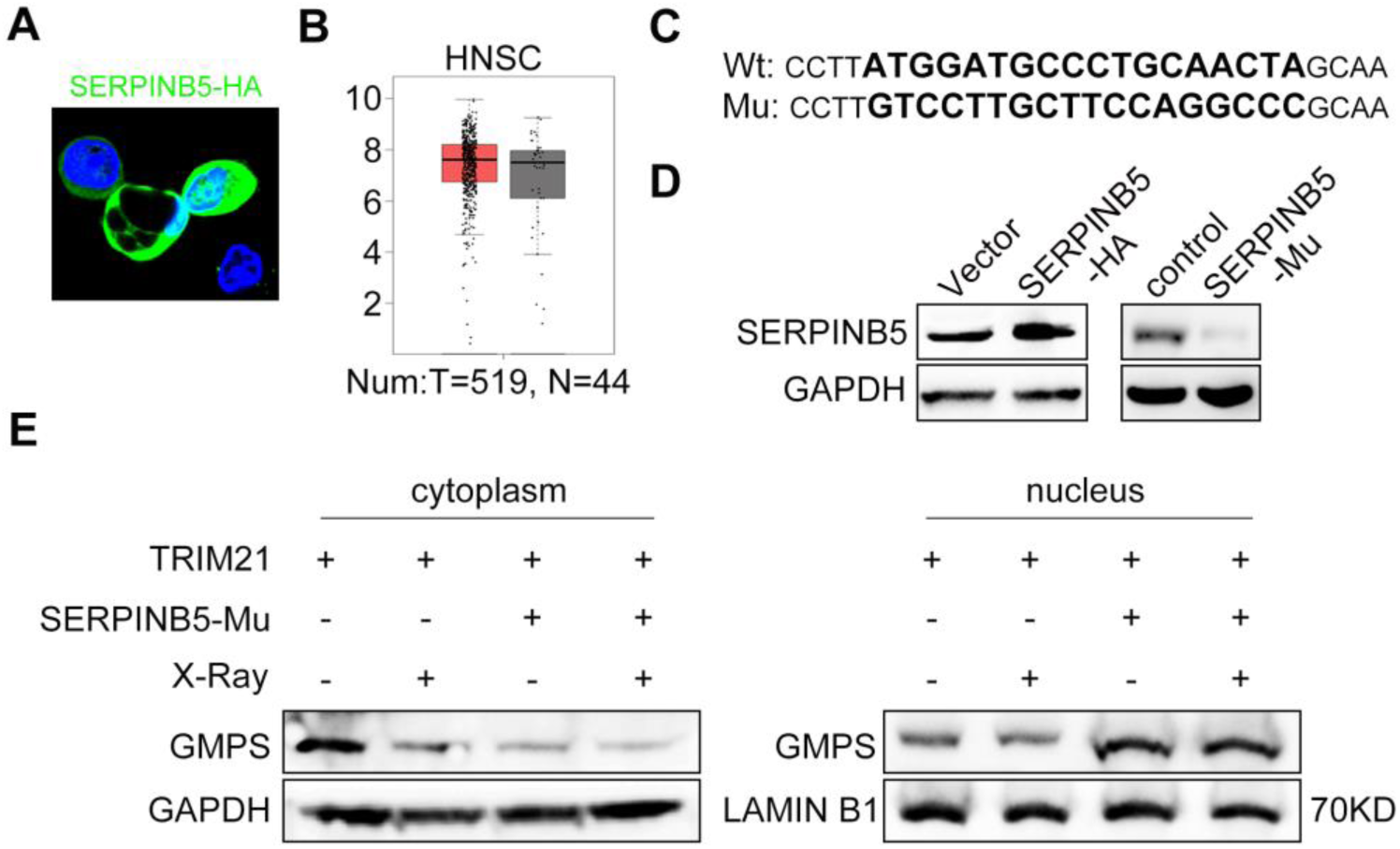
SERPINB5 is essential for TRIM21-mediated GMPS repression. A Immunofluorescence staining of SERPINB5–HA in HONE1 cells. B *SERPINB5* mRNA expression levels in HNSC in TCGA dataset. T: Tumor, N: normal. C CRISPR-mediated *SERPINB5* knockout NPC cells. Bolded, larger typeface indicates the mutated sequences. D SERPINB5 expression in *SERPINB5* knockout HONE1 cells. E GMPS expression in cytoplasm (left) or nucleus (right) of HONE1 cells with *TRIM21* overexpression or *SERPINB5* LOF. Data information: Wt, wild-type; Mu, mutant.

**Supplementary Figure 3.**
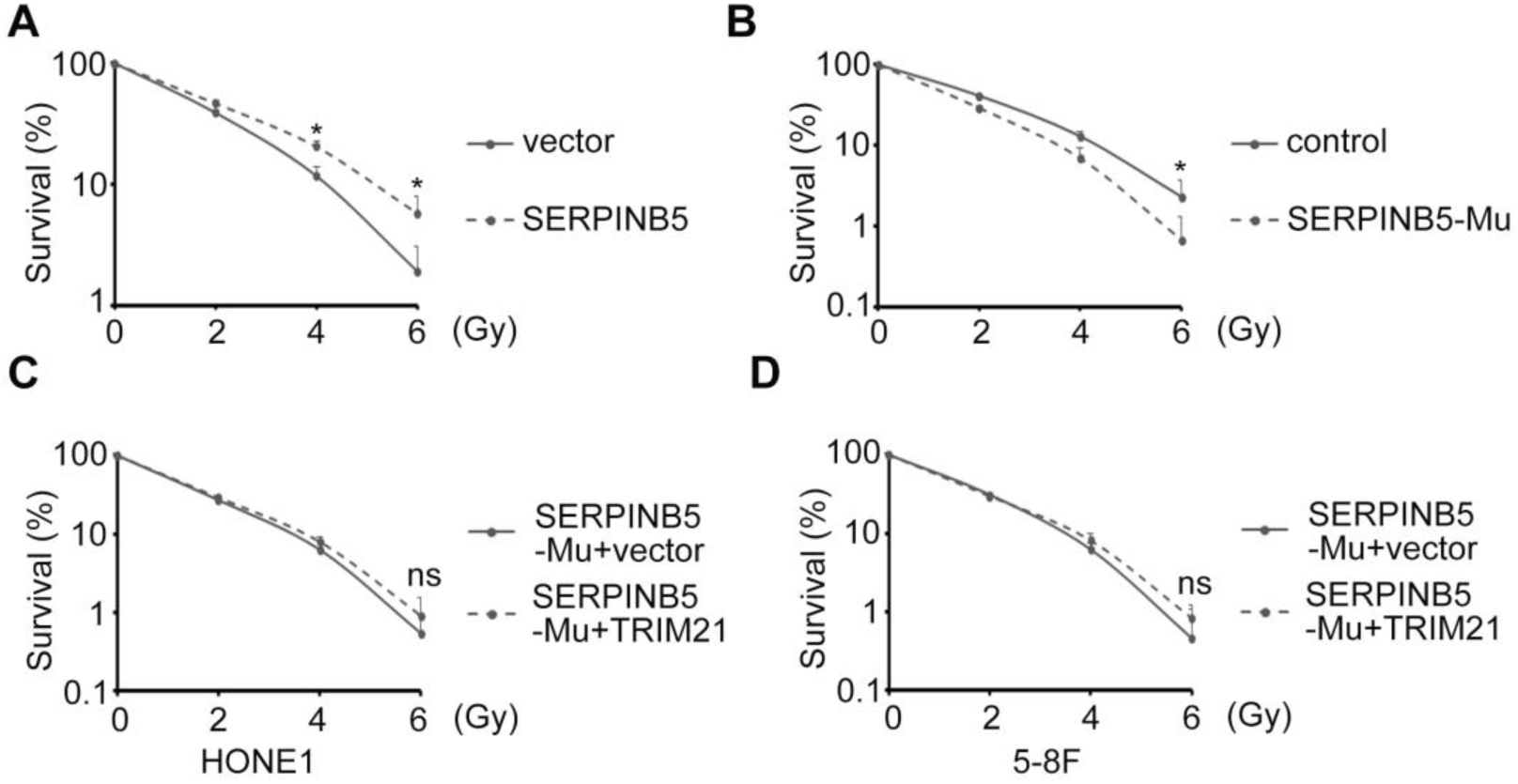
SERPINB5 is essential for TRIM21-mediated NPC cell survival after radiation. A, B The survival rates of HONE1 cells with *SERPINB5* GOF (A) or LOF (B) after radiation. C, D The survival rates of HONE1 (C) or 5-8F (D) cells with *SERPINB5* knockout and *TRIM21* GOF. Data information: Mu, mutant.

**Supplementary Figure 4.**
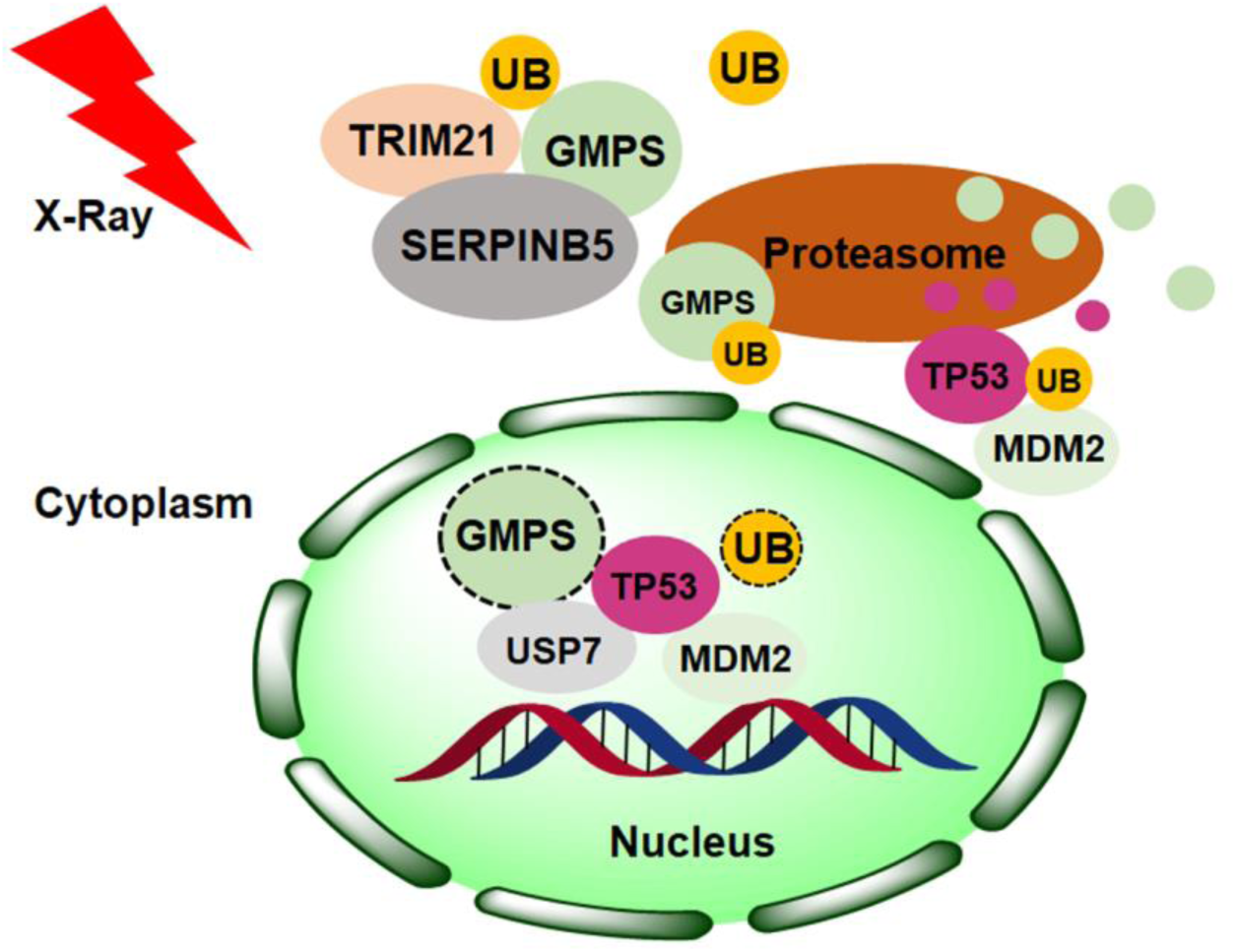
The working model of TRIM21–SERPINB5-mediated GMPS–TP53 repression in NPC cells after X-ray radiation. Data information: UB, ubiquitin

